# Endopeptidase regulation as a novel function of the Zur-dependent zinc starvation response

**DOI:** 10.1101/398487

**Authors:** Shannon G. Murphy, Laura Alvarez, Myfanwy C. Adams, Shuning Liu, Joshua S. Chappie, Felipe Cava, Tobias Dörr

## Abstract

The cell wall is a strong, yet flexible, meshwork of peptidoglycan (PG) that gives a bacterium structural integrity. To accommodate a growing cell, the wall is remodeled by both PG synthesis and degradation. *Vibrio cholerae* encodes a group of three nearly identical zinc-dependent endopeptidases (EPs) that hydrolyze PG to facilitate cell growth. Two of these (*shyA* and *shyC*) are housekeeping genes and form a synthetic lethal pair, while the third (*shyB*) is not expressed under standard laboratory conditions. To investigate the role of ShyB, we conducted a transposon screen to identify mutations that activate *shyB* transcription. We found that *shyB* is induced as part of the Zur-mediated zinc starvation response, a mode of regulation not previously reported for cell wall lytic enzymes. *In vivo*, ShyB alone was sufficient to sustain cell growth in low-zinc environments. *In vitro*, ShyB retained its D,D-endopeptidase activity against purified sacculi in the presence of the metal chelator EDTA at a concentration that inhibits ShyA and ShyC. This suggests that ShyB can substitute for the other EPs during zinc starvation, a condition that pathogens encounter while infecting a human host. Our survey of transcriptomic data from diverse bacteria identified other candidate Zur-regulated endopeptidases, suggesting that this adaptation to zinc starvation is conserved in other Gram-negative bacteria.

**Importance:** The human host sequesters zinc and other essential metals in order to restrict growth of potentially harmful bacteria. In response, invading bacteria express a set of genes enabling them to cope with zinc starvation. In *Vibrio cholerae*, the causative agent of the diarrheal disease cholera, we have identified a novel member of this zinc starvation response: a cell wall hydrolase that retains function in low-zinc environments and is conditionally essential for cell growth. Other human pathogens contain homologs that appear to be under similar regulatory control. These findings are significant because they represent, to our knowledge, the first evidence that zinc homeostasis influences cell wall turnover. Anti-infective therapies commonly target the bacterial cell wall and, therefore, an improved understanding of how the cell wall adapts to host-induced zinc starvation could lead to new antibiotic development. Such therapeutic interventions are required to combat the rising threat of drug resistant infections.

## Introduction

The cell wall provides a bacterium with structural integrity and serves as a protective layer guarding against a wide range of environmental insults. Due to its importance for bacterial survival, the cell wall is a powerful and long-standing target for antibiotics (1). The wall is composed primarily of peptidoglycan (PG), a polymer of β-(1,4) linked N-acetylglucosamine (NAG) and N-acetylmuramic acid (NAM) sugar strands (2) (**Fig. 1A**). NAM peptide side chains are cross-linked to peptides on adjacent strands, enabling the PG to assemble into a meshlike structure called the sacculus (3). In Gram-negative bacteria, the sacculus is a single PG layer that is sandwiched between an inner and an outer membrane (4). This thin wall must be rigid enough to maintain cell shape and to contain high intracellular pressures (3, 5). However, the wall must also be flexible enough to accommodate cell elongation, cell division, and the insertion of *trans*-envelope protein complexes (6). This requirement for both rigidity and flexibility necessitates continuous remodeling of the cell wall, which is accomplished by a delicate interplay between PG synthesis and degradation. Inhibition or dysregulation of process can cause growth cessation or cell lysis, rendering the mechanisms of cell wall turnover an attractive target for new antibiotic development (7, 8).

**Fig 1.**
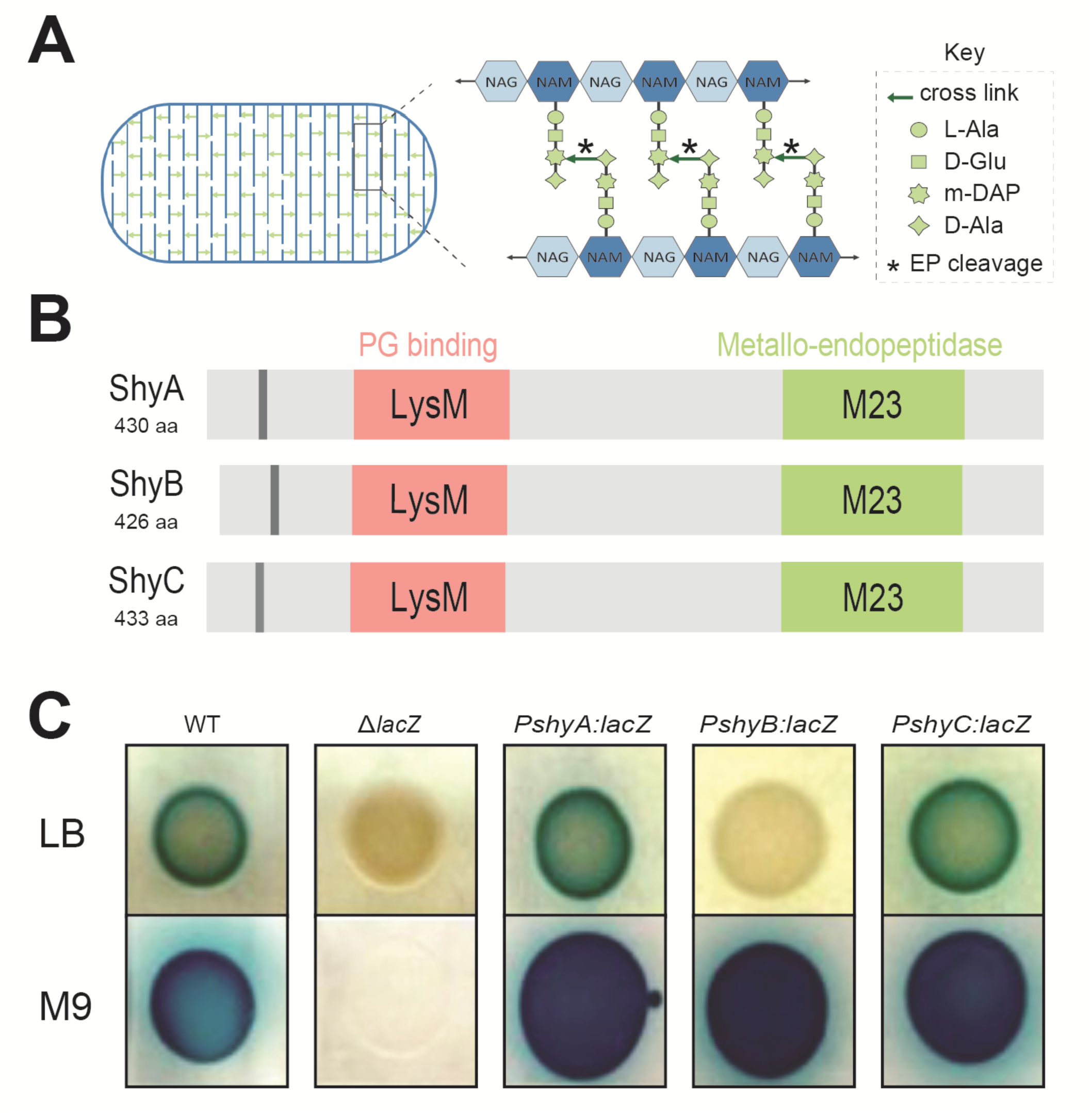
*shyB* is a LysM/M23 endopeptidase that is transcribed in minimal medium. **(A)** Model of the peptidoglycan sacculus indicating EP cleavage sites **(B)** The *V. cholerae* genome encodes three endopeptidases (ShyA, ShyB, ShyC) possessing a hydrophobic region (gray), a PG binding domain (LysM, pink), and metallo-endopeptidase domain (M23, green). Protein domains were annotated using UniProt (65). **(C)** *lacZ* transcriptional reporters for each endopeptidase spotted onto LB (top row) and M9 minimal (bottom row) agar containing X-gal. A blue colony color indicates that the promoter is actively transcribed. Wild-type (WT) and Δ*lacZ* strain are included as positive and negative controls, respectively.

PG synthesis is mediated by Penicillin Binding Proteins (PBPs, the targets for beta-lactam antibiotics) and SEDS proteins (9). These proteins collectively catalyze cell wall synthesis through two main reactions: transglycosylation (TG) to elongate the sugar backbone and transpeptidation (TP) to crosslink the peptide stems of adjacent strands (2). Cell wall turnover is mediated by “autolysins”, a collective term for diverse and often redundant enzymes (amidases, carboxypeptidases, lytic transglycosylases and endopeptidases) that are able to cleave PG at almost any chemical bond (6). Endopeptidases (EPs), for example, hydrolyze the peptide crosslinks that covalently link adjacent PG strands, effectively reversing the TP reaction. EPs are crucial for cell elongation in both Gram-positive and Gram-negative rod-shaped bacteria (10-12), presumably because they create gaps in the PG meshwork to allow for the insertion of new cell wall material. Consistent with this proposed role, EP overexpression promotes aPBP activity in *Escherichia coli*, likely through the generation of initiation sites for PG synthesis (13).

While EPs are essential for growth, they are also main drivers of PG degradation after inhibition of PBPs (14, 15). Thus, EP activity must be tightly controlled under normal growth conditions. EPs in two divergent bacterial species (*E. coli* and *Pseudomonas aeruginosa*) are proteolytically degraded to adapt to conditions that require changes in PG cleavage activity (16, 17), such as the transition into stationary phase. In *Bacillus subtilis*, EP expression is regulated by growth-phase dependent sigma factors (18-21). However, it is not known how EP expression is modified in response to specific environmental stresses.

In this study, we investigate the role of specialized EPs in *V. cholerae*, the causative agent of the diarrheal disease cholera. *V. cholera* encodes three nearly identical EPs that are homologous to the well-characterized D,D-endopeptidase MepM in *E. coli* (10). Each EP contains a LysM domain that likely binds PG (22) and a Zn^2+^-dependent M23 catalytic domain that hydrolyzes peptide cross links (23) (**Fig. 1B**). We previously showed that two of these (ShyA and ShyC) are housekeeping EPs that are collectively essential for growth (12). The gene encoding the third EP, *shyB*, is not transcribed under standard laboratory conditions (LB medium) and thus little is known about its biological function. To elucidate the role of ShyB, we conducted a transposon screen to identify mutations that promote *shyB* expression in LB. We found that *shyB* is induced by zinc starvation and, unlike the other two M23 EPs, ShyB enzymatic activity is resistant to treatment with the metal chelator EDTA. These data suggest that ShyB acts as an alternative EP to ensure proper PG maintenance under zinc limiting conditions. Importantly, this represents the first characterization of an autolysin that is controlled by Zur-mediated zinc homeostasis and provides insight into how other Gram-negative pathogens might adapt to zinc-starvation when colonizing a human host.

## Results

### *shyB* is repressed in LB, but transcribed in minimal medium

The hydrolytic activity of autolysins needs to be carefully controlled to maintain cell wall integrity. We therefore considered it likely that specialized autolysins are transcriptionally regulated and only induced when required. To test this, we examined expression patterns of the LysM/M23 endopeptidases using *lacZ* transcriptional fusions. We first compared promoter activity on LB and M9 agar, as our previous work showed that a Δ*shyB* mutation exacerbates a Δ*shyA* growth defect in M9 minimal medium (12). The *P*_*shyA*_*:lacZ* and *P*_*shyC*_*:lacZ* reporters generated a blue colony color on both LB and M9 minimal agar (**Fig. 1C**), meaning that these promoters are actively transcribed on both media. This is consistent with ShyA and ShyC’s role as housekeeping EPs (12). In contrast, *P*_*shyB*_*:lacZ* yielded blue colonies on M9 minimal medium only, indicating that the *shyB* promoter is induced in M9 but repressed in LB.

### *shyB* is induced by zinc starvation

To elucidate the specific growth conditions that favor *shyB* expression, we sought to identify the genetic factors controlling *shyB* transcription. To this end, we subjected the transcriptional reporter strain to Himar1 mariner transposon mutagenesis and screened for P_*shyB*_*:lacZ* induction (blue colonies) on LB agar. After two independent rounds of mutagenesis (50,000 total colonies), the screen yielded 26 blue colored insertion mutants. These were divided into two distinct classes according to colony color: 12 dark blue and 14 light blue colonies. Strikingly, arbitrary PCR (24) mapped all 26 transposon insertions to two chromosomal loci involved in zinc homeostasis: *vc0378/zur* (dark blue colonies) and *vc2081-2083*/*znuABC* (light blue colonies) (**Fig. 2A**). Zur is a fur-family transcriptional regulator and the central repressor in the zinc starvation response (25). In zinc-rich conditions, Zur and its Zn^2+^ corepressor bind to promoters containing a “Zur box” and block transcription (26). In low-zinc conditions, Zur dissociates from promoters to induce the zinc starvation response (27). This regulon includes genes encoding zinc uptake systems (i.e. *znuABC, zrgABCDE*) (28) and zinc-independent paralogs that replace proteins that ordinarily require zinc for function (i.e. ribosomal proteins) (29). The Zur-controlled *znuABC* locus encodes *V. cholerae’*s high affinity zinc uptake system (28). To validate the transposon hits, we constructed clean deletions of *zur* and *znuA* in the P_*shyB*_*:lacZ* reporter strain. Deletion of either gene resulted in activation of the *shyB* promoter on LB agar, and *shyB* repression was restored by expressing the respective genes *in trans* (**Fig. S1**). Thus, *shyB* is induced under conditions that are expected to either mimic (*zur* inactivation) or impose (*znuA(BC*) inactivation) zinc starvation.

**Fig 2.**
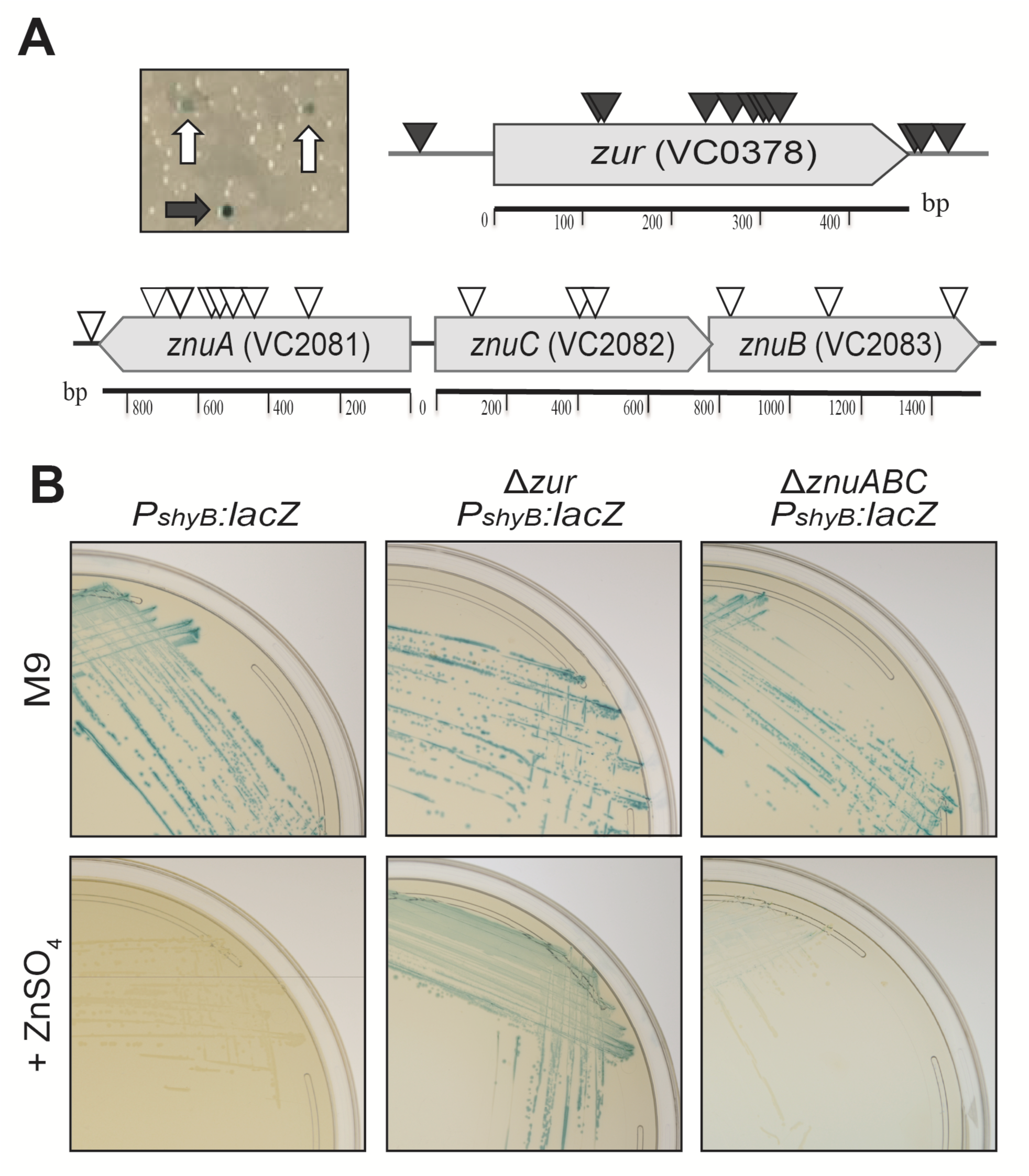
*shyB* transcription is regulated by zinc homeostasis. **(A)** The *shyB* transcriptional reporter (*lacZ::P*_*shyB*_*:lacZ*) was mutagenized with a Himar1 mariner transposon and screened for *shyB* induction (blue colonies) on LB agar containing X-gal and selective antibiotics (see Methods). Representative dark blue (black arrow) and light blue (white arrows) colonies are shown. Approximate Tn insertion sites identified by arbitrary PCR are shown (triangles). **(B)** The *shyB* transcriptional reporter in a wild-type, Δ*zur*, or Δ*znuABC* background were grown on M9 X-gal agar without (top row) or with (bottom row) 10 µM ZnSO_4_.

If zinc starvation is the factor inducing *shyB* expression in M9, we would expect the P_*shyB*_*:lacZ* reporter to be repressed by external zinc addition. Indeed, supplementing M9 with 10 µM of ZnSO_4_ was sufficient to turn off the *shyB* promoter in a wild-type (WT) background (**Fig. 2B**), whereas repression could not be achieved by adding in other transition metals (iron and manganese) (**Fig. S2**). In a Δ*zur* background, the *shyB* promoter remained active even when M9 was supplemented with exogenous zinc (**Fig. 2B**), indicating that Zur is required for P_*shyB*_ repression. We also found that zinc supplementation somewhat repressed the *shyB* promoter in Δ*znuA*, suggesting that *V. cholerae* can uptake zinc even in the absence of its primary transporter. Indeed, *V. cholerae* encodes a second, lower affinity zinc acquisition system (*zrgABCDE*) to maintain zinc homeostasis (28).

### Zur directly binds the *shyB* promoter

Given Zur’s well-defined role as a transcriptional regulator (26) and its requirement for *P*_*shyB*_ repression in zinc-rich media, we hypothesized that Zur directly binds the *shyB* promoter. To test this, we retrieved a Zur box sequence logo built from 62 known regulatory targets in Vibrionaceae (30, 31) and aligned it with the *shyB* promoter region. This alignment identified a highly conserved Zur box characterized by an inverted, AT-rich repeat (**Fig. 3A**). We used 5’-RACE to locate the *shyB* transcriptional start site (tss) and found that the putative Zur box overlaps both the −10 region and tss. A bound Zur/Zn^2+^ complex at this position likely prevents RNA polymerase binding and thereby prevents transcription (32).

**Fig 3.**
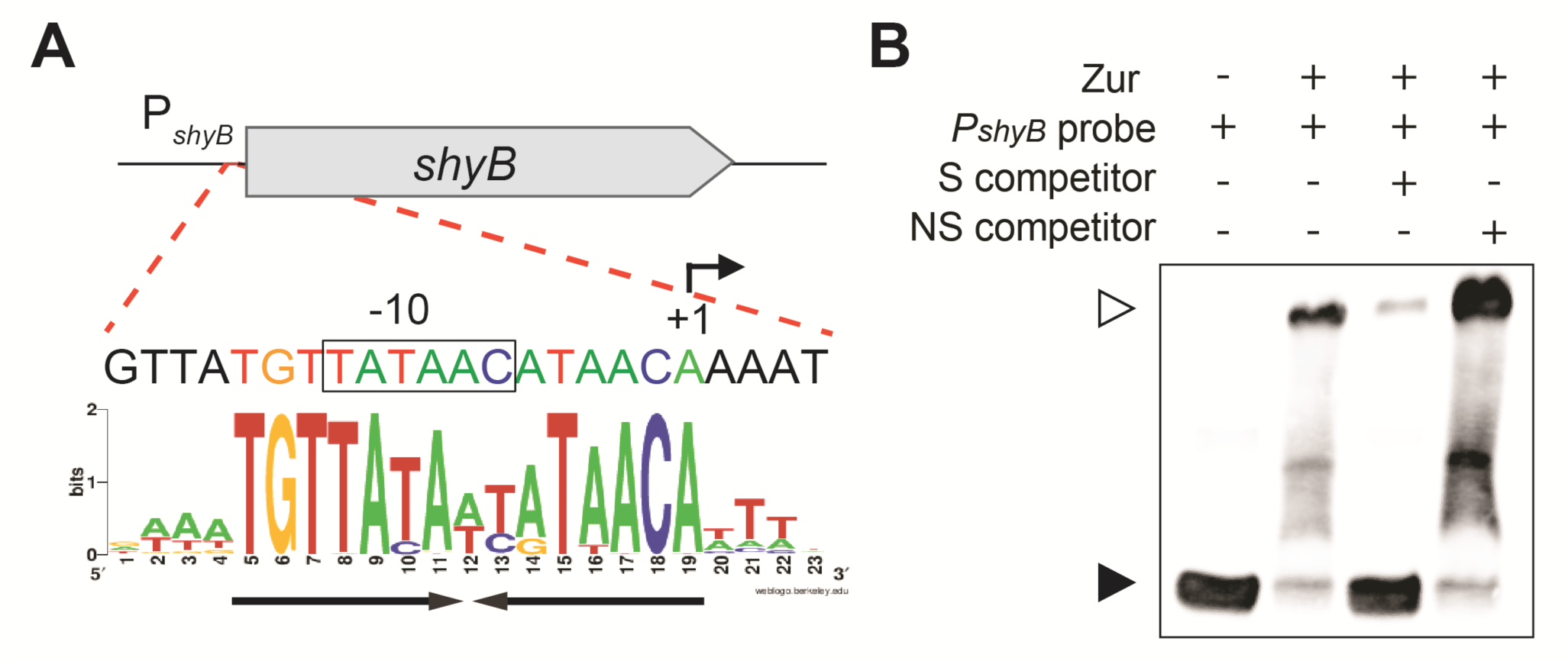
Zur directly binds the *shyB* promoter. **(A)** The *shyB* promoter, annotated with a 5’-RACE transcription start site (+1) and putative −10 region (box), was aligned with a *Vibrio* Zur sequence logo (30, 31). The inverted AT-rich repeat in the putative Zur-box is underlined with black arrows. **(B)** A chemiluminescent probe containing the putative *shyB* Zur box was incubated with purified Zur in the presence of ZnCl_2_ (5 µM). Zur binding specificity was tested by adding 100-fold molar excess of unlabeled specific (S, Lane 3) or non-specific (NS, Lane 4) competitor DNA. Samples were electrophoresed on a 6% DNA retardation gel to separate unbound (black arrow) and bound probe (white arrow).

To determine if Zur binds the *shyB* promoter *in vitro*, we incubated purified Zur with a labeled DNA probe encoding the *P*_*shyB*_ Zur box. Binding was assessed in the presence of ZnCl_2_ using an electrophoretic mobility shift assay (EMSA). As evident by a band shift, Zur formed a complex with the P_*shyB*_ DNA *in vitro* (**Fig. 3B, Lanes 1-2**). To examine DNA binding specificity, a 100-fold molar excess of unlabeled specific (S) or non-specific (NS) competitor DNA was included in the binding reaction. The S competitor, which carries an identical sequence as the labeled probe, effectively sequestered Zur and increased the amount of unbound, labeled probe (**Lane 3**). Meanwhile, the NS competitor was ineffective at binding Zur (**Lane 4**). These data indicate that the *shyB* promoter contains an authentic Zur box and we conclude that *shyB* is a novel member of the Zur regulon.

### ShyB supports growth in chelated medium

As *shyB* is part of the Zur-mediated zinc starvation response, we hypothesized that ShyB endopeptidase activity supports cell growth when zinc availability is low. To induce zinc starvation and robustly derepress the Zur regulon, *V. cholerae* strains were grown in M9 minimal medium supplemented with TPEN (N,N,N′,N′-Tetrakis(2-pyridylmethyl)ethylenediamine), an intracellular metal chelator with high affinity for zinc (33). As expected from our genetic analysis, TPEN addition resulted in the production of ShyB protein, which could be reversed by adding zinc (**Fig. S3**).

We first tested whether native *shyB* could restore Δ*shyAC* growth under zinc-starvation conditions. *shyA* and *shyC* deletions were generated in a parent strain expressing an IPTG-inducible copy of *shy*A (*lacZ*::P_tac_:*shyA*), as these genes are conditionally essential in rich media (12). In the absence of IPTG, we found that chelation with either TPEN or EDTA (a more general divalent metal ion chelator), induced growth of Δ*shyAC*, but not in the mutant that additionally lacked *shyB* (**Fig. 4A**; **Fig. S4**). As expected, chelation-dependent growth of Δ*shyAC* could be suppressed by adding zinc (**Fig. 4B**; **Fig. S4**). These data suggest that induction of *shyB* alone is sufficient to sustain *V. cholerae* growth, and synthetic lethality of *shyA* and *shyC* is due to the lack of *shyB* expression under laboratory growth conditions. Consistent with this interpretation, we were able to generate a Δ*shyAC* knockout in a Δ*zur* background (**Fig. S5**) or in a strain exogenously overexpressing *shyB* (**Fig. S6**).

**Fig 4.**
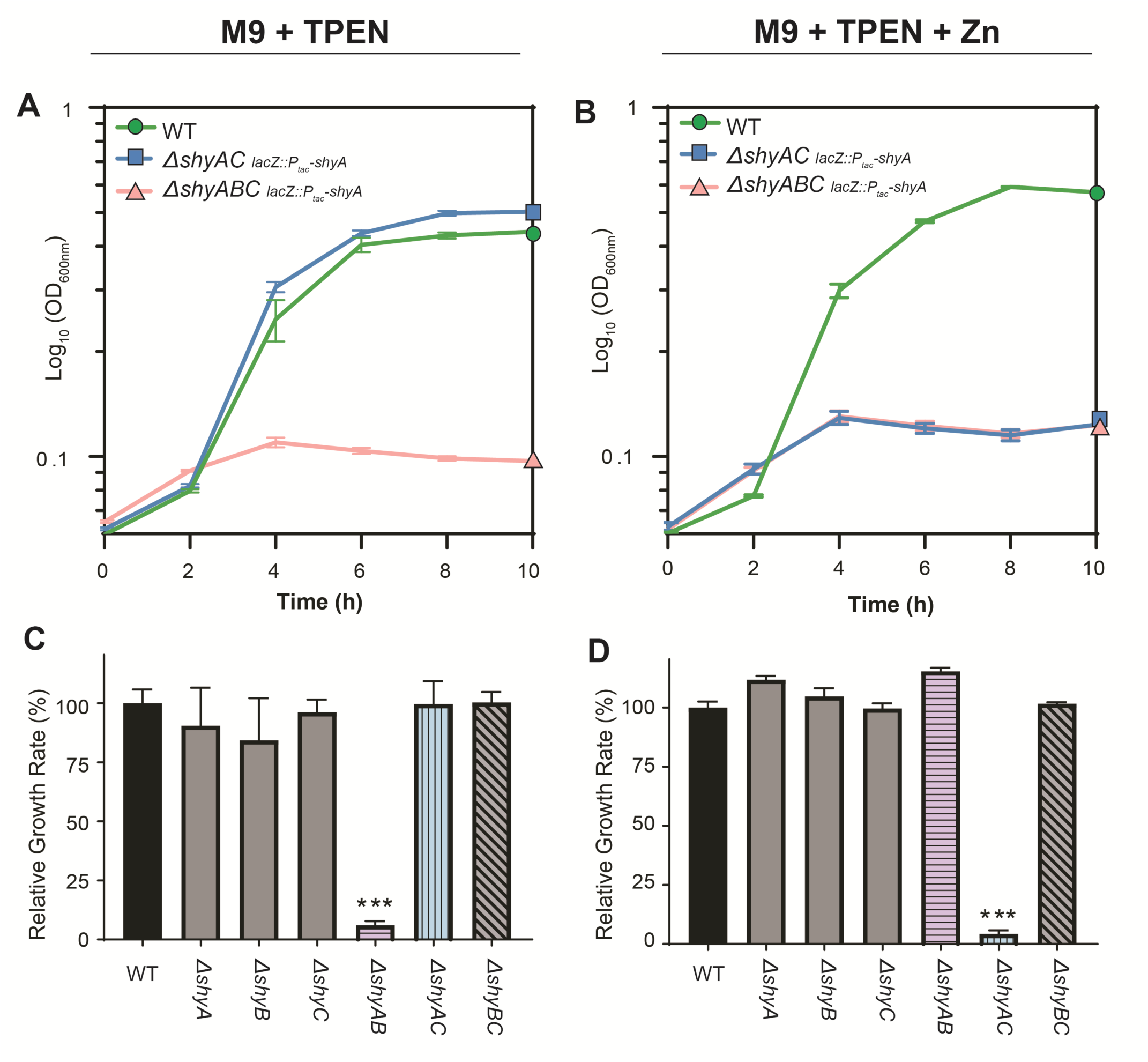
*shyB* supports cell growth in chelated medium and is conditionally essential in a Δ*shyA* mutant. Mid-exponential cultures of the indicated *V. cholerae* mutants were washed to remove IPTG before being diluted 1:100 into M9 glucose containing streptomycin plus TPEN (250 nM) in the absence (A,C) or presence (B,D) of ZnSO_4_ (1 µM). Growth of each strain was monitored by optical density (OD600) in a Bioscreen C 100-well plate. Error bars report standard error of the mean (SEM) for three independent biological replicates. **(A-B)** Log-transformed growth curves are shown for WT (green circle), Δ*shyAC lacZ::P*_*tac*_*:shyA* (blue square), and Δ*shyABC lacZ::P*_*tac*_*:shyA* (red triangle). **(C-D)** In a similar growth experiment, growth rates of WT (solid black), single mutants (solid gray) and double mutants (striped) were calculated from exponential phase and normalized to the average WT growth rate (%). Statistical difference relative to WT was assessed using two-way analysis of variance (ANOVA) followed by Dunnett’s multiple comparison test (***, p-value < 0.001).

A Δ*shyB* mutant alone did not exhibit a significant growth defect in M9 TPEN (**Fig. 4C**); however, autolysins often need to be deleted in combination to elicit a substantial phenotype (11). We therefore generated all possible combinations of LysM/M23 endopeptidase deletions to broadly dissect the relevance of zinc concentrations for EP activity. Of these, the Δ*shyAB* double mutant failed to grow in the presence of TPEN (**Fig. 4C**). This indicates that ShyC, the only essential LysM/M23 EP in the Δ*shyAB* mutant, cannot support growth in zinc-starved media. In contrast, only the Δ*shyAC* mutant failed to grow in zinc-replete medium and this can be explained by Zur-mediated *shyB* repression (**Fig. 4D**). This tradeoff in synthetic lethality partners tentatively suggests that ShyB may function as a replacement for ShyC during zinc starvation. ShyC protein levels, as measured by Western Blot, were not reduced in the presence of TPEN, ruling out the possibility that Δ*shyAB* lethality reflects transcriptional downregulation or degradation of ShyC (**Fig S3**). Rather, these observations suggest that ShyC activity is more sensitive to zinc-chelation than the other EPs. Alternatively, TPEN might induce changes in PG architecture that make it resistant to cleavage by ShyC.

### ShyB is an EDTA-resistant D,D-endopeptidase *in vitro*

ShyB is predicted to be a D,D-endopeptidase but biochemical evidence is lacking. Thus, we measured the *in vitro* hydrolytic activity of each EP against *V. cholerae* sacculi. Each protein was recombinantly purified without the hydrophobic signal sequence or transmembrane domain (ShyA_Δ1-35_, ShyB_Δ1-34_, and ShyC_Δ1-33_) to increase stability *in vitro*. As a negative control, we purified ShyB with a mutation (H370A) in the active site that is expected to abolish activity. EPs were incubated with purified sacculi (see Methods for details) and the soluble PG fragments released by digestion were separated using high pressure liquid chromatography (HPLC) and quantified by spectrophotometry. As predicted, all three enzymes, but not the H370A mutant, hydrolyzed *V. cholerae* sacculi and generated soluble PG fragments (**Fig. 5A**; **Fig. S7**). Sacculi digestion with ShyA and ShyC resulted in a similar profile of PG fragments, indicating similar hydrolytic activity *in vitro*. In contrast, the ShyB chromatogram showed more peaks with shorter retention times. These observations suggest that ShyB further processes the sacculi into smaller fragments. Consistent with this, ShyB was able to further process PG pre-digested with ShyA or ShyC, while these EPs only slightly modified ShyB-digested PG (**Fig. S8**). To determine which muropeptides remain in the insoluble pellet after EP digestion, muramidase was used to digest the PG sugar backbone (β 1→4 linkages). The resulting soluble products produced a single peak, corresponding to a M4 monomer. This indicates that all three LysM/M23 EPs exhibit D,D-endopeptidase activity *in vitro* (**Fig. 5B**).

**Fig 5.**
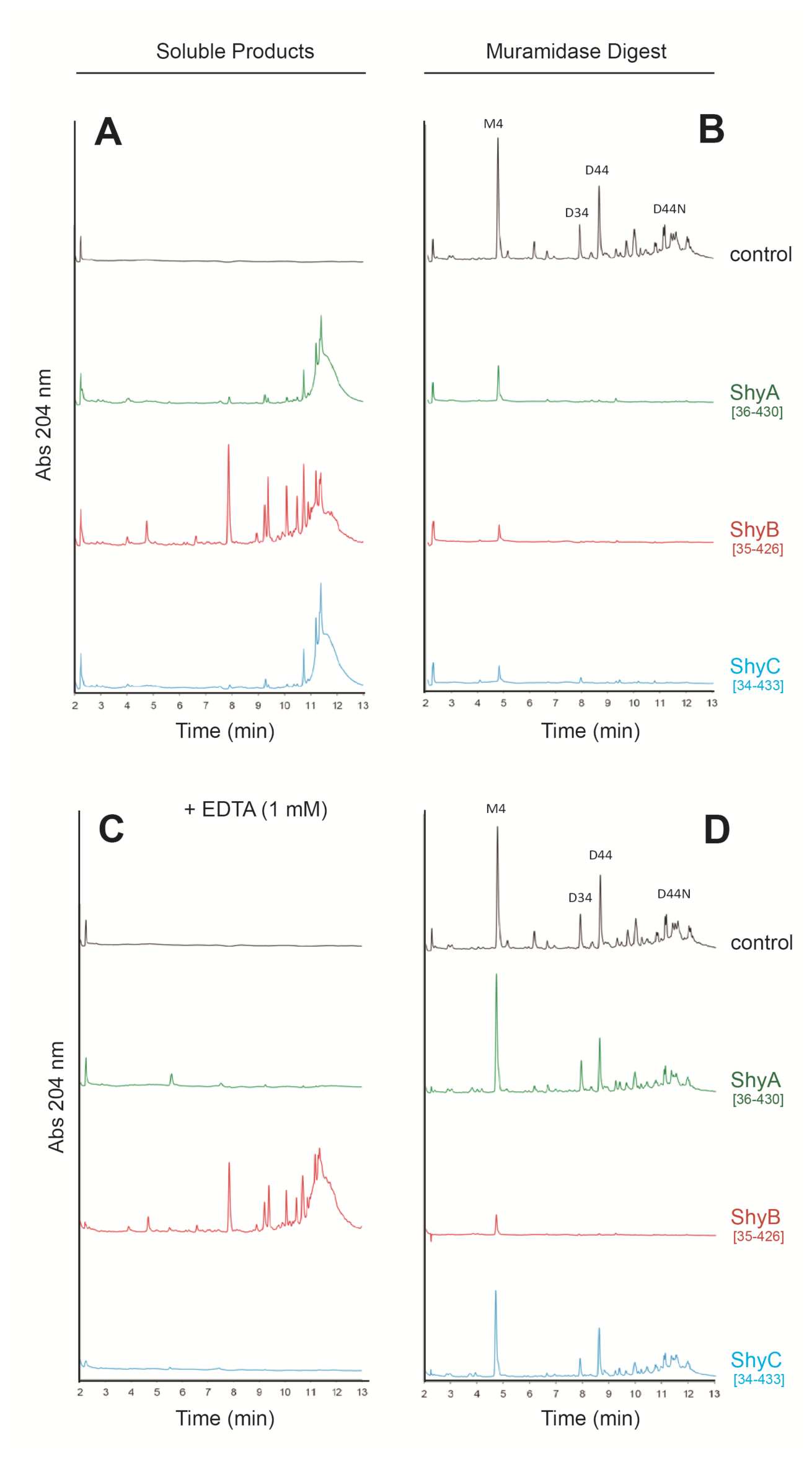
ShyB retains endopeptidase activity in EDTA. *V. cholerae* sacculi was digested with 10 µg of purified ShyA, ShyB, or ShyC for 16 h at 37°C in the absence **(A)** or presence **(C)** of 1mM EDTA. The soluble products released by digested sacculi were separated by size via HPLC and quantified by absorbance (204 nm). (**B, D**) The remaining insoluble pellet was digested with muramidase and soluble products were separated by HPLC.

M23 domains require a coordinated zinc ion to carry out PG hydrolysis (23). However, based on its regulation by Zur, we hypothesized that ShyB evolved to function in zinc-limited environments. To test this, we repeated the *in vitro* PG hydrolysis assays under metal-limited conditions by using the divalent cation chelator EDTA. Strikingly, ShyB retained its activity in the presence of EDTA (1 mM), while ShyA and ShyC activity was completely abolished (**Fig. 5C-d**). This is consistent with results previously obtained for ShyA (12). These data suggest that ShyB has a high affinity for, or can function without, divalent cations like zinc.

### ShyB localizes to the division septum

Endopeptidases often differ in their cellular localization and this is an important determinant of EP function *in vivo* (34). We previously reported that ShyA, a sidewall hydrolase, remains diffuse throughout the periplasm, while ShyC localizes to the septum (midcell) during division (12). To investigate the relative role of ShyB, we constructed a functional ShyBmsfGFP translational fusion (**Fig. S9**) and visualized its localization using epifluorescence microscopy. ShyBmsfGFP strongly localized to the septum as cells prepared to divide. (**Fig. 6**). Septal localization suggests that ShyB and ShyC are involved in cell division. However, neither the Δ*shyB* nor Δ*shyC* mutant, either alone or in combination, have a division defect and we thus do not know the significance of endopeptidase activity at the septum in *V. cholerae.*

**Fig 6.**
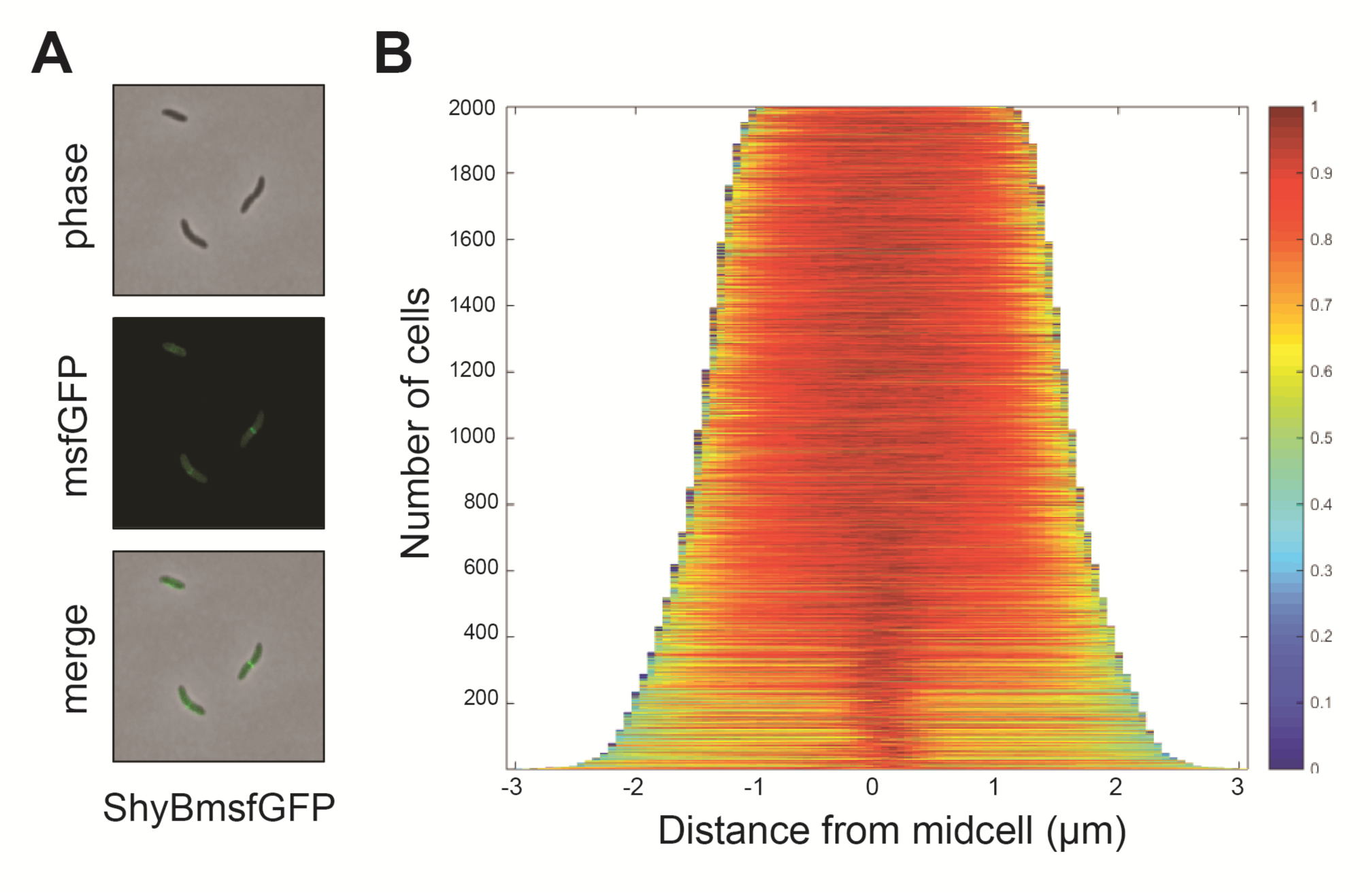
ShyBmsfGFP localizes to the midcell during division. *V. cholerae* expressing a C-terminal fluorescent fusion (*lacZ::P*_*tac*_*-shyBmsfGFP*) was grown overnight in M9 + IPTG (200 µM). (A) The ShyBmsfGFP fusion was imaged on an agarose patch (0.8% agarose in M9 minimal medium) with 300 ms exposure at 490 nm. (B) A heat map showing intensity of fluorescent signal as a function of distance from the midcell (“demograph”) was generated from over 1,800 cells in Oufti (64).

### Zur-controlled endopeptidases are widespread in divergent bacteria

Zur-controlled EPs appear to be widespread in *Vibrionaceae*. Using BLAST homology searches, we have identified isolates from 30 different non-cholera *Vibrio* species that contain a ShyB homolog with a Zur box upstream of the gene encoding it (**Table S1**) (35). To assess the significance of zinc homeostasis for EP regulation more broadly, we surveyed published microarray and RNAseq datasets from diverse bacteria for differential EP expression (36-45). *Yersinia pestis* CO92, the causative agent of plague, encodes a ShyB/MepM homolog (YPO2062) that is significantly up-regulated in a Δ*zur* mutant (36). YPO2062 does not contain its own Zur box, but is adjacent to *znuA* and may thus be co-transcribed as part of the same operon (**Fig. 7**). Similarly, *mepM* (b1856) is located adjacent to the *znu* operon in laboratory (K12 MG1655) and pathogenic *E. coli* (Enterohemorrhagic O157:H7 and Enteropathogenic O127:H6). Two independent microarray studies in *E. coli*, one of which was validated by qPCR, showed that this EP was transcriptionally upregulated in response to zinc starvation (44, 45). These data suggest that MepM and its homologs are Zur-regulated EPs. Notably, this *znu*/EP arrangement is conserved in many other Gram-negative pathogens, including *Salmonella typhimurium* (STM1890), *Enterobacter cloacae* (ECL_0-1442), and *Klebsiella pneumoniae* (KPK_1913). Lastly, *A. baumannii*, an important nosocomial pathogen, possesses two M23 endopeptidases differentially transcribed in a Δ*zur* mutant. A1S_3329 is up-regulated and A1S_0820 is down-regulated compared to a wild-type strain (37), suggesting that these EPs are also under zinc starvation control. Collectively, these data suggest that zinc homeostasis and cell wall turnover are linked in a wide array of Gram-negative bacteria.

**Fig 7.**
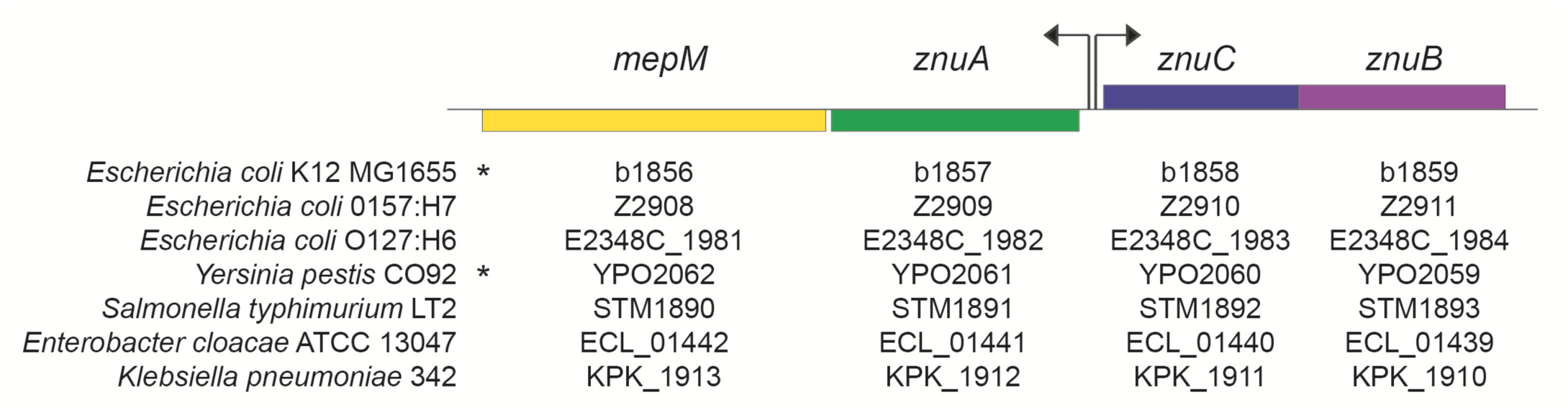
ShyB/MepM homologs are adjacent to the Zur-controlled *znu* operon in many Gram-negative pathogens. Gene neighborhood alignments generated by Prokaryotic Sequence Homology Analysis Tool (PSAT) from 7 different Gram-negative bacteria (66). Arrows indicate the approximate location of the bidirectional promoter and site of Zur-binding in the *znu* operon. Asterisks indicate that co-transcription of *znu* and the downstream M23 endopeptidase is supported by transcriptomic data.

## Discussion

### Highly redundant endopeptidases support cell growth

Endopeptidase activity is essential for cell growth in both Gram-negative and Gram-positive bacteria, supporting the long-standing hypothesis that autolysins create space in the PG meshwork for the insertion of new cell wall material (8). As with other autolysins, EPs are highly redundant but exhibit slight differences in cellular localization (i.e. septal or sidewall) (12, 20, 46), substrate specificity (10,47) and relative abundance during each growth phase (11, 46). Our previous work in *V. cholerae* identified three LysM/M23 zinc metallo-endopeptidases: two (ShyA and ShyC) are housekeeping enzymes that are conditionally essential for growth, while the third (ShyB) is not expressed under standard laboratory conditions (12). In this study, we define *shyB* as a new member of the Zur regulon and demonstrate that ShyB can replace the other EPs *in vivo* when zinc concentrations are limiting. This is a novel mechanism for regulating autolysins and establishes a link between two essential processes: cell wall turnover and metal ion homeostasis.

### Zinc availability affects the expression and activity of cell wall hydrolases

Zur represses *shyB* transcription in zinc-rich growth conditions. This is consistent with our initial observation that the *shyB* promoter is active on M9 and repressed on LB agar. These respective media differ markedly in terms of zinc content; M9 contains no added zinc while LB naturally contains high levels of zinc ions (∼12.2 µM) (48). We found that adding zinc (10 µM) to M9 represses the *shyB* promoter and hence the zinc starvation response. As a cautionary note, this suggests that *V. cholerae* is starved for zinc in M9, a complication not usually considered when interpreting results obtained in this medium.

Based on its membership in the Zur regulon, it is likely that ShyB evolved to function in low-zinc environments. Indeed, ShyB endopeptidase activity is resistant to high EDTA concentrations *in vitro*. In an apparent contradiction, the ShyB crystal structure models a zinc ion in the active site (49). It is possible that ShyB has a higher affinity for zinc than the other EPs, but we cannot yet exclude the possibility that ShyB utilizes other metal cofactors. ShyA appears to have an intermediate ability to function in low zinc environments; we found that ShyA can support cell growth in the absence of the other two EPs in TPEN-treated medium (like ShyB), yet EDTA inhibited its activity *in vitro*. This observation is likely a consequence of the high EDTA concentrations (1 mM) used in the biochemical assays, which do not permit wild type *V. cholerae* growth. The ability to sustain growth in the presence of TPEN, however, indicates that ShyA function is less affected by metal starvation than ShyC.

Since ShyA functions in chelated medium, we tentatively hypothesize that *shyB* is derepressed to compensate for a loss of ShyC activity. This model is supported by localization data and ShyC’s sensitivity to chelating conditions, both *in vivo* and *in vitro*, that induce *shyB* expression. We did not observe any defects in Δ*shyB*, Δ*shyC*, or Δ*shyBC* mutants; however, septal EP deletion causes division defects (i.e. filamentation) in other bacteria (20). It is thus possible that the role of septal EPs in *V. cholerae* is more prominent under conditions not yet assayed. In our experiments, diffuse ShyA might be present at sufficient concentrations at the septum to alleviate any obvious division defects.

### Bacteria encounter zinc starvation while infecting a host

Proteins that retain function in low-zinc conditions likely play important roles in pathogenesis as bacteria encounter zinc-starvation inside the human host (50). Vertebrates and other organisms sequester metals to restrict the growth of potentially harmful bacterial, a defense strategy referred to as “nutritional immunity” (50). In response, bacteria employ zinc-starvation responses to maintain essential cellular processes (51). Zinc importers (*znuABC* and *zrgABCDE*), for example, are critical for host colonization and infection in *V. cholerae* (28), *A. baumannii* (52), pathogenic *E. coli* (53, 54), *Salmonella enterica* (55), and others (56). It is tempting to speculate that ShyB, a rather unusual addition to the Zur regulon, supports PG remodeling in a zinc-depleted host environment. Consistent with this idea, *shyB* is located on a mobile genomic island (VSP-II) that is strongly associated with the current (seventh) Cholera pandemic (57). The current pandemic strain emerged in the 1960’s and, owing to its higher spread capability, replaced its pandemic predecessors (58). This suggests that VSP-II (and thus possibly ShyB) conferred a fitness advantage to pathogenic *V. cholerae*.

Importantly, Zur-controlled M23 endopeptidases do not appear to be confined to *V. cholerae*. Diverse bacteria, including notable human pathogens, possess a conserved *shyB*/*mepM*/*yebA* homolog adjacent to the Zur-controlled *znu* operon. Transcriptomic data from both *Y. pestis* and *E. coli* support the prediction that this EP is upregulated along with the zinc importer. The conservation of zinc-regulated EPs in divergent Gram-negative pathogens suggests that there may be a widely conserved mechanism for maintaining cell wall homeostasis in low zinc environments. Importantly, this may confer an important adaptation to host-induced zinc starvation. These findings in *V. cholerae* will inform future investigations examining the interplay between cell wall turnover and zinc homeostasis.

#### Experimental Procedures

### Bacterial growth conditions

Cells were grown by shaking (200 rpm) at 37°C in 5 mL of LB medium unless otherwise indicated. M9 minimal medium with glucose (0.4%) was prepared with ultrapure Mili-Q water to minimize zinc contamination. When appropriate, antibiotics were used at the following concentrations: streptomycin (200 µg mL^-1^), ampicillin (100 µg mL^-1^), and kanamycin (50 µg mL^-1^). IPTG (200 µM) was added to all liquid and solid media if required to sustain *V. cholerae* growth. X-gal (40 µg mL^-1^) was added to plates for blue-white screening.

### Plasmid and strain construction

All genes were PCR amplified from *V. cholerae* El Tor N16961 genomic DNA. Plasmids were built using isothermal assembly (59) with the oligonucleotides summarized in **Table S2**. The suicide vector pCVD442 was used to make gene deletions via homologous recombination (60); 700 bp regions flanking the gene of interest were amplified for Δ*zur* (SM89/90 + SM91/92), Δ*znuA* (SM107/108 + SM109/110), and Δ*znuABC* (SM93/94, SM95/96) and assembled into XbaI digested pCVD442. Endopeptidase deletion constructs were built as described previously (12). Chromosomal delivery vectors (pJL-1 and pTD101) were used to insert genes via double cross-over into native *lacZ*. To construct the *shyB* transcriptional reporter, 500 bp upstream of *shyB* were amplified (SM1/2) and assembled into NheI-digested pAM325 to yield a *P*_*shyB*_*:lacZ* fusion. This fusion was amplified (SM3/4) and cloned into StuI-digested pJL-1 (61). To complement gene deletions, *zur* (SM99/100) and *znuA* (SM113/114) were cloned into SmaI-digested pBAD: a chloramphenicol resistant, arabinose-inducible plasmid. To construct the ShyBmsfGFP C-terminal translational fusion, *shyB* (SM181/63) and *msfGFP* (SM65/66) were amplified with an overhang encoding a 10 amino acid flexible linker (gctggctccgctgctggttctggcgaattc). These fragments were assembled into SmaI-digested pTD101, which positions the fusion under an IPTG-inducible promoter. In a similar manner, pTD101(*shyB*) was constructed with SM181/182 and pTD100(*shyA*) was built as previously described (12). An additional chromosomal delivery vector (pSGM100) was built for crossover into VC0817. *<u>shyB</u>* (SM141/SM55) was placed under arabinose-inducible control by cloning into SmaI-digested pSGM100. All assemblies were initially transformed into *E. coli* DH5a λpir and then into SM10 λpir for conjugation into *V. cholerae.*

All strains are derivatives of *V. cholerae* El Tor N16961 (WT). To conjugate plasmids into *V. cholerae*, SM10 λpir donor strains carrying pCVD442, pTD101, PJL-1, or pSGM100 plasmids were grown in LB/ampicillin. Recipient *V. cholerae* strains were grown overnight in LB/streptomycin. Stationary phase cells were pelleted by centrifugation (6,500 rpm for 3 min) and washed with fresh LB to remove antibiotics. Equal ratios of donor and recipient (100µL:100 µL) were mixed and spotted onto LB agar plates. After a 4-hour incubation at 37°C, cells were streaked onto LB containing streptomycin and ampicillin to select for cross-over recipients. Colonies were purified and cured through two rounds of purification on salt free sucrose (10%) agar with streptomycin. Insertions into native *lacZ* (via pJL-1, pTD101) were identified by blue-white colony screening on X-gal plates. Gene deletions (via pCVD442) were checked via PCR screening with the following primers: Δ*shyA* (TD503/504), Δ*shyB* (SM30/31), Δ*shyC* (TD701/702), Δ*zur* (SM122/123), Δ*znuA* (SM119/120), and Δ*znuABC* (SM119/121).

### Transposon mutagenesis and arbitrary PCR

The *shyB* transcriptional reporter was mutagenized with Himar1 mariner transposons, which were delivered via conjugation by an SM10 λpir donor strain carrying pSC189 (62). The recipient and donor were grown overnight in LB/streptomycin and LB/ampicillin, respectively. Stationary phase cells were pelleted by centrifugation (6,500 rpm for 3 min) and washed with fresh LB to remove antibiotics. Equal ratios of donor and recipient (500 µL:500 µL) were mixed and spotted onto 0.45 µm filter disks adhered to pre-warmed LB plates. After a 4-hour incubation at 37°C, cells were harvested by aseptically transferring the filter disks into conical tubes and vortexing in fresh LB. The cells were spread onto LB agar containing streptomycin to kill the donor strain, kanamycin to select for transposon mutants, and X-gal to allow for blue-white colony screening. Plates were incubated at 30°C overnight followed by two days at room temperature. To identify the transposon insertion site, purified colonies were lysed via boiling and used directly as a DNA template for arbitrary PCR. As described elsewhere, this technique amplifies the DNA sequence adjacent to the transposon insertion site through successive rounds of PCR (24). Amplicons were Sanger sequenced and high quality sequencing regions were aligned to the *N16961* genome using BLAST (35).

### 5’ rapid amplification of cDNA ends

The *shyB* transcription start site was identified with 5’ rapid amplification of cDNA ends (5’ RACE). To obtain a *shyB* transcript, Δ*zur* was grown in LB at 37°C until cells reached mid-exponential phase (OD_600_ = 0.5) and RNA was extracted using Trizol and acid:phenol chloroform (Ambion). DNA contamination was removed through two RQ1 DNase (Promega) treatments and additional acid:phenol chloroform extractions. cDNA synthesis was performed with MultiScribe reverse transcriptase (ThermoFisher) and a *shyB* specific primer (SM270). cDNA was column purified and treated with terminal transferase (New England BioLabs) to add a homopolymeric cytosine tail to the 3’ end. The cDNA was amplified through two rounds of touchdown PCR with a second gene-specific primer (SM271) and the Anchored Abridged Primer (ThermoFisher). The PCR product was Sanger sequenced using primer SM271.

### Electrophoretic mobility shift assay

The LightShift Chemiluminescent EMSA kit (ThermoFisher) was used to detect Zur-promoter binding. 41 bp complimentary oligos (SM264/265) containing the putative *shyB* Zur box, with and without a 5’ biotin label, were annealed according to commercial instructions (Integrated DNA Technologies). 20 µL binding reactions contained buffer, Poly dI-dC (50 ng µL^-1^), ZnCl_2_ (5 µM), labeled probe (1 pmol), and purified Zur (600 nM). Unlabeled specific or non-specific competitor oligos were added in 100-fold molar excess. Reactions were incubated on ice for 1 hour, electrophoresed on a 6% DNA retardation gel (100 V, 40 min), and wet transferred to a Biodyne B membrane (100 V, 30 min) (ThermoFisher) in a cold room. The membrane was developed using chemiluminescence according to the manufacturer’s instructions and imaged using a Bio-Rad ChemiDoc MP imaging system.

### Protein expression and purification

DNA encoding N-terminally truncated LysM/M23 endopeptidases (ShyA_Δ1-35_, ShyB_Δ1-34_, and ShyC_Δ1-33_) and full length Zur was PCR amplified from genomic DNA, while template for the ShyB H370A mutation was commercially synthesized (Integrated DNA Technologies). Shy constructs were cloned into pCAV4, and Zur into pCAV6, both modified T7 expression vectors that introduce an N-terminal 6xHis-NusA tag (pCAV4) or 6xHis-MBP tag (pCAV6) followed by a Hrv3C protease site upstream of the inserted sequence. Constructs were transformed into BL21(DE3) cells, grown at 37°C in Terrific Broth supplemented with carbenicillin (100 mg mL^-1^) to an OD_600_ of 0.8-1.0, and then induced with IPTG (0.3 mM) overnight at 19°C. ZnCl_2_ (50 µM) was added during Zur induction. Cells were harvested via centrifugation, washed with nickel loading buffer (NLB) (20 mM HEPES pH 7.5, 500 mM NaCl, 30 mM imidazole, 5% glycerol (v:v), 5 mM β –Mercaptoethanol), pelleted in 500mL aliquots, and stored at −80°C.

Pellets were thawed at 37°C and resuspended in NLB supplemented with PMSF (10 mM), DNAse (5 mg), MgCl_2_ (5 mM), lysozyme (10 mg mL^-1^), and one tenth of a complete protease inhibitor cocktail tablet (Roche). All buffers used in Zur purification were supplemented with ZnCl_2_ (1 µM). Cell suspensions were rotated at 4°C, lysed via sonication, centrifuged, and the supernatant was syringe filtered using a 0.45 µM filter. Clarified samples were loaded onto a NiSO_4_ charged 5 mL HiTrap chelating column (GE Life Sciences), and eluted using an imidazole gradient from 30 mM to 1M. Hrv3C protease was added to the pooled fractions and dialyzed overnight into cation exchange loading buffer (20 mM HEPES pH 7.5, 50 mM NaCl, 1 mM EDTA, 5% glycerol (v:v), 1 mM DTT). Cleaved Shy proteins were loaded onto a 5 mL HiTrap SP HP column and cleaved Zur was loaded onto a 5mL HiTrap Heparin HP column (GE Life Sciences). All constructs were eluted along a NaCl gradient from 50mM to 1M. Fractions were concentrated and injected onto a Superdex 75 16/600 equilibrated in Size Exclusion Chromotography buffer (20 mM HEPES pH7.5, 150 mM KCl, 1 mM DTT). Zur dimers coeluted with MBP on the sizing column and were subsequently incubated with amylose resin (New England BioLabs) at 4°C and Zur was collected from a gravity column. Final purified protein concentrations were determined by SDS-PAGE and densitometry compared against BSA standards: ShyA, 5.72 mg mL^-1^; ShyB, 5.72 mg mL^-1^; ShyB H320A, 2.35 mg mL^-1^; ShyC, 17.93 mg mL^-1^; Zur, 0.31 mg mL^-1^.

### Sacculi digestion assay

Peptidoglycan from stationary phase *V. cholerae* cells was extracted and purified via SDS boiling and muramidase digestion (63). 10 µL of sacculi and 10 µg of enzyme were mixed in 50 µL buffered solution (50 mM Tris-HCl pH 7.5, 100 mM NaCl) in the absence or presence of 1 mM EDTA. Digestions were incubated for 16 h at 37°C. Soluble products were harvested and the remaining pellet was further digested with muramidase. All soluble products were reduced with sodium borohydride, their pH adjusted, and injected into a Waters UPLC system (Waters, Massachusetts, USA) equipped with an ACQUITY UPLC BEH C18 Column, 130 Å, 1.7 µm, 2.1 mm × 150 mm (Waters) and a dual wavelength absorbance detector. Eluted fragments were separated at 45°C using a linear gradient from buffer A [formic acid 0.1% (v/v)] to buffer B [formic acid 0.1% (v/v), acetonitrile 40% (v/v)] in a 12 min run with a 0.175 ml min^-1^ flow, and detected at 204 nm.

### Growth curve analysis

Strains were grown overnight in LB/streptomycin with IPTG. Cells were washed in 1X phosphate buffered solution (PBS) and subcultured 1:10 into M9 glucose plus IPTG. After 2 hours shaking at 37°C, cells were washed and subcultured 1:100 into M9 glucose containing combinations of TPEN (250 nM), ZNSO4 (1 µM), and IPTG (200 µM). The growth of each 200 µL culture in a 100-well plate was monitored by optical density (OD_600_) on a Bioscreen C plate reader (Growth Curves America).

### Microscopy and image analysis

Cells were imaged on an agarose patch (0.8% agarose in M9 minimal medium) using a Leica DMi8 inverted microscope. To image the ShyBmsfGFP fusion, cells were exposed to 490 nm for 300 ms. Image analysis, including cell selection and subpixel quantification of fluorescent signal as a function distance from the midcell, was performed in Oufti (64).

## Acknowledgements

We thank all the members of the Doerr and Chappie Labs for helpful discussions and assistance with this work. We thank members of the John Helmann Lab for lending their expertise and reagents. We thank the faculty, staff, and students at the Weill Institute for Cell and Molecular Biology (WICMB) for their support. Research in the Doerr Lab is supported by start-up funds from Cornell University. Research in the Chappie Lab is supported by the National Institute of Health (NIH). Research in the Cava lab is supported by MIMS, the Knut and Alice Wallenberg Foundation (KAW), the Swedish Research Council and the Kempe Foundation.

---

**Fig S1.** Δ***zur* and** Δ***znuABC* deletions induce the *shyB* promoter on LB agar.** Clean deletions of Δ*zur* and Δ*znuA* in the *P*_*shyB*_*:lacZ* transcriptional reporter were complemented with an arabinose-inducible (pBAD) plasmid carrying the respective gene *in trans*. Strains were plated onto LB agar containing x-gal (40 µg mL^-1^), chloramphenicol (10 µg mL^-1^), and arabinose (0.2%). Plates were incubated overnight and then at room temperature for 2 days.

**Fig S2. *shyB* promoter is repressed by exogenous zinc, but not by other transition metals.** The *P*_*shyB*_*:lacZ* transcriptional reporter was plated on M9 X-gal agar containing 10 µM of ZnSO_4_, FeSO_4_, or MnCl_2_. Plates were incubated overnight and then at room temperature for 2 days.

**Fig S3. Western Blot of ShyB and ShyC protein levels in high and low-zinc media**. N16961 strains encoding ShyB ΔlysM::6His-FLAG or ShyC:6His-FLAG were grown in M9 glucose (0.4%) with added TPEN (250 nM) or TPEN plus ZnSO4 (1 µM). Cells were harvested at mid-log (OD600 = 0.4) and lysed via SDS boiling and sonication. Western blot was performed using standard techniques. Blots were developed using a mouse anti-FLAG F1804 primary antibody (Sigma Aldrich) and Goat anti-Mouse IR CW800 secondary antibody (LI-COR Biosciences). Blots were imaged using a Lycor Odyssey CLx imager.

**Fig S4. EDTA-induced *shyB* expression restores growth to Δ*shyAC***. Wt (green), Δ*shyAC lacZ::P*_*tac*_*-shyA* (blue), and Δ*shyABC lacZ::P*_*tac*_*-shyA* (red) strains were grown in M9 glucose (0.4%) containing **(A)** EDTA (30 µM) (solid lines) or **(B)** EDTA plus ZnSO_4_ (60 µM) (dashed lines). Growth of each 200 µL culture was measured by optical density (600 nm) in a Bioscreen C 100-well plate. Error bars report standard error of the mean (SEM) for three biologically independent replicates.

**Fig S5. *zur* deletion restores growth to Δ*shyAC* in LB medium.** Overnight cultures (grown in LB/streptomycin at 37°C) were subcultured 1:100 into fresh media and grown at 37°C until mid-log phase. Δ*zur lacZ::P*_*tac*_*-zur* (blue) and Δ*zur* Δ*shyAC lacZ::P*_*tac*_*-zur* (red) were diluted 1:100 into LB (solid lines) or in LB plus IPTG (200 µM) (dashed lines). Growth of each strain was monitored by optical density (OD600) in a Bioscreen C 100-well plate. Error bars report standard error of the mean (SEM) for three biologically independent replicates.

**Fig S6. Inducible *shyB* expression rescues growth of Δ*shyABC.*** Strains were grown overnight in LB/streptomycin plus IPTG (200uM) at 37°C. Cells were washed, subcultured 1:10 into M9 glucose (0.4%), and grown at 37°C for 2 hours. Wt (dotted lines) and Δ*shyABC lacZ::P*_*tac*_*-shyA* vc1807*::P*_*ara*_*-shyB* (solid lines) strains were diluted 1:100 in M9 glucose (0.4%) (orange), with 200 µM IPTG (green) or with 0.2% arabinose (black). Growth of each 200 µL culture was measured by optical density (600 nm) in a Bioscreen C 100-well plate. Error bars report standard error of the mean (SEM) for three biologically independent replicates.

**Fig S7. A point mutation in the ShyB active site abolishes endopeptidase activity *in vitro***. Purified ShyB, and ShyB H370A were incubated with purified *V. cholerae* sacculi for 16 h at 37°C. **(A)** The soluble products released by digested were separated by HPLC and quantified by absorbance (204 nm). **(B)** The remaining pellet was digested with muramidase and the soluble products were separated by HPLC and quantified by absorbance.

**Fig S8. Sequential digestion of *V. cholerae* sacculi by Shy endopeptidases.** 10 µg of purified (**A**) ShyA, (**B**) ShyB, and (**C)** ShyC were incubated with *V. cholerae* sacculi for 16 h at 37°C, followed by secondary digestion a different endopeptidase. The soluble products released by digested sacculi were separated by size via HPLC and quantified by absorbance (204 nm).

**Fig S9. ShyBmsfGFP translational fusion rescues growth of Δ*shyAB* in TPEN-chelated medium.** Mid-exponential cultures of WT (blue), Δ*shyAB* (red), Δ*shyAB lacZ::P*_*tac*_*-shyB* (green), and Δ*shyAB lacZ::P*_*tac*_*-shyB* (orange, purple) were washed and subcultured 1:100 into M9 containing TPEN (250 nM) with and without IPTG (200 µM). Growth of each culture at 37°C was measured by optical density (600 nm). Error bars report standard error of the mean (SEM) for three biologically independent replicates.

**Table S1. Summary of ShyB homologs that contain an upstream, canonical Zur box.**

**Table S2. Summary of oligonucleotides used in this study.**

